# Ribavirin shows antiviral activity against SARS-CoV-2 and downregulates the activity of TMPRSS2 and the expression of ACE2 In Vitro

**DOI:** 10.1101/2020.12.04.410092

**Authors:** Mehmet Altay Unal, Ceylan Verda Bitirim, Gokce Yagmur Summak, Sidar Bereketoglu, Inci Cevher Zeytin, Omur Bul, Cansu Gurcan, Dunya Aydos, Ezgi Goksoy, Ebru Kocakaya, Zeynep Eran, Merve Murat, Nil Demir, Julia Somers, Emek Demir, Hasan Nazir, Sibel Aysil Ozkan, Aykut Ozkul, Alpay Azap, Acelya Yilmazer, Kamil Can Akcali

## Abstract

Ribavirin is a guanosine analog and has a broad-spectrum antiviral activity against RNA viruses. Based on this, we aimed to show the anti-SARS-CoV-2 activity of this drug molecule via in vitro, in silico and molecular techniques. Ribavirin showed antiviral activity in Vero E6 cells following SARS-CoV-2 infection. In silico analysis suggested that Ribarivin has a broad-spectrum impact on Vero E6 cells. According to the detailed molecular techniques, Ribavirin was shown to decrease TMPRSS2 expression both at mRNA and protein level 48 hours after treatment. The suppressive effect of Ribavirin in ACE2 protein expression was shown to be dependent on cell types. Finally, proteolytic activity assays showed that Ribavirin also showed an inhibitory effect on TMPRSS2 enzyme. As a conclusion, Ribavirin is a potential antiviral drug for the treatment against SARS-CoV-2, and it interferes with the effect of TMPRSS2 and ACE2 expression.

## Introduction

The coronavirus diseases 2019 (COVID-19) pandemic represents the hardest global public health threat since the influenza outbreak in 1918 and has rapidly resulted in significant health and economic burden worldwide [1]. Pre-clinical and clinical trials for the treatment of COVID-19 have been ongoing in various centers across the World. Currently, there are 2271 clinical trials which involves therapeutic or preventative interventions registered at www.clinicaltrials.gov. Among these trials, different therapeutic approaches have been studies including antiviral, antimalarial, immunomodulator and cell/plasma based therapies [1]. However, results of pre-clinical and clinical studies may vary between each other and there is still no definite therapeutic strategy.

Drug repurposing is a method preferred by researchers to introduce new therapeutics, reduce the time and financial burden of developing molecules or drugs [2]. Ribavirin, one of the FDA-approved antiviral drugs used for the last four decades against RNA viruses, could be a potential candidate to be used for drug repurposing against COVID-19. Ribavirin (1-β-D-ribofuranosyl-1,2,4-triazole-3-carboxamide) with broad-spectrum antiviral properties, first synthesized in 1972 [3], has been used mostly to treat hepatitis C virus (HCV) infections in humans. Although ribavirin is used almost exclusively in hepatitis C virus infections in clinical practice, it has an antiviral effect against different viruses [4]. Furthermore, ribavirin has suitable formulations for inhalation, or intravenous administration and oral use. Due to this feature, it has been also preferred in the treatment of different viral infections. It is well-known that ribavirin molecule has different antiviral mechanisms of action against different viruses [5]. In one these mechanisms, ribavirin monophosphate can potently inhibit guanosine derivatives (eg guanosine triphosphate [GTP]) and reduce nucleotide residues resulting in antiviral effect, especially in studies with respiratory syncytial virus (RSV), yellow fever virus and paramyxovirus [6-9]. Secondly, the main intracellular metabolite of ribavirin is ribavirin triphosphate (RTP) which can act through polymerase inhibition. RTP can bind competitively with molecules such as adenosine triphosphate (ATP) or GTP and inducing the inhibition of polymerases. This mechanism of action has been observed in studies conducted with influenza virus, reovirus and vesicular stomatitis virus [4, 10, 11]. As a final mechanism of action; ribavirin molecule can bind with translation initiation factor 4E (eIF4E), preventing the initiation of translation, or interacting with the enzymes responsible for RNA cap synthesis, preventing the initiation of translation. This mechanism has been reported via in silico molecular docking analysis for the SARS-CoV-2 virus and Lassa fever [12]. Ribavirin is also used in Crimean-Congo haemorrhagic fever, coronavirus, South American haemorrhagic fevers, hantavirus and adenovirus infections [4]. As stated above, ribavirin exerts antiviral effects in different viruses via exerting different mechanisms. Both its ease of use and its ability to be used against different viruses; suggest that ribavirin is a promising potent and broad-spectrum antiviral drug. For this reason, in this study, the antiviral effect of ribavirin against SARS-CoV-2 has been studies in detail via in silico, in vitro and molecular biology analysis.

## Materials & Methods

### Cells and viruses

The Vero E6 (African green monkey, kidney epithelial) cells were incubated in a humidified atmosphere at 37 °C and 5% CO_2_. They were incubated in high glucose Dulbecco’s’ modified Eagle medium (DMEM) (Capricorn) supplemented with 10% fetal bovine serum (Biolegend), 100 U/mL of penicillin (Lonza) and 0.1 mg/mL of streptomycin (Lonza). Human colonadeno carcinoma Caco-2 cells were maintained in DMEM high glucose supplemented with 10% fetal bovine serum, 100 μg/mL streptomycin, 100 U/mL penicillin, and 1% non-essential amino acids. To subcultivate, cells were incubated with trypsin/EDTA solution at 37 °C until cells detached followed by washing with phosphate buffered saline (PBS). They were centrifuged at 900 x g for 3 minutes. Then they were seeded at a concentration around 0.23×10^6^ cell/mL. To stock early passage cells, the growth media including 10% DMSO (Sigma, cell culture grade) was used.

A local isolate of SARS-CoV-2 (Ank1) was used in this study. Viruses were propagated in Vero E6 cells by using DMEM media containing 10% FBS and 1% antibiotics. All virus-related experiments were performed at Biosafety Level 3 laboratories.

### Viral Infection

Vero E6 cells were seeded onto 96-well plates prior to infection experiment. Cells were infected with SARS-CoV-2-Ank1 isolate at 100DKID_50_/mL dose. Cells were incubated at 37°C incubator 1 h to allow viral penetration. After 1h, Ribavirin was added into the culture media by serially diluting the main stock (100 µM) via 5-fold dilutions. Four wells were used for each experimental condition. Plates were monitored for the presence of cytopathic effect specific (CPE) to SARS-CoV-2-Ank1.

### In silico calculation

Calculations are done with AutoDock-Vina software[1]. BIOVIA-Discovery Studio [2], and UCSF Chimera v1.14 [3] software are used for the imaging. Swiss ADME (http://www.swissadme.ch/) online tool is used for detailed ADME-T calculations. Also, molecular docking calculations were made to model ACE2-COVID-19 interaction and all regions of COVID-19. Proteins used in the calculations are obtained from the protein data bank (PDB) (https://www.rcsb.org/).

### MTT Assay

Evaluation of toxic doses of Ribavirin was analyzed by MTT assay. Vero E6 and Caco-2 cells were plated on 96-well plates at a concentration of 10.000 cells/well and incubated 24 hours. Then, the cells were treated with 10, 25, 200, and 750 µM Ribavirin for 24 and 48 hours. ddH2O was used as solvent. Cell proliferation was assessed by the MTT assay (Milipore (CT02). Absorption was measured at 570 nm using Epoch™ Microplate Spectrophotometer. Then the OD values were analyzed as percentage of cell viability relative to the control.

### Flow Cytometry

TMPRSS2 expression of Ribavirin treated cells and control cells was evaluated by flow cytometry (NovoCyte Flow Cytometry). Vero E6 and Caco2 cells were harvested with Trypsin-EDTA (2X) and single cell suspensions were stained using antibody against TMPRSS2 (Santa Cruze, sc-515727) after fixation (4% Paraformaldehyde, 15 min RT) and permeabilization (INTRA, 15 min RT) process. All analysis was performed using Novo Express 1.3.0 Software (ACEA Biosciences, Inc).

### mRNA expression by qRT-PCR

To show alteration of mRNA expression levels of TMPSS2 and ACE2 after drug treatment, quantitative real-time PCR (qRT-PCR) was performed. Total RNA isolation was done using total RNA isolation solution (GeneAll® Ribo Ex^™^, 301-001). 1 µg total RNA was used to cDNA synthesis (TONBO Biosciences, 31-5300-0100R). qRT-PCR studies were performed using TONBO Biosciences CYBER Fast™ qPCR Hi-ROX Master Mix. Gene expression was normalized to the expressions of GAPDH and analyzed with the 2-ΔΔCT method as fold change compared to solvent control.

### Western Blotting

ACE2 protein level was evaluated by western blotting after drug treatment, cells were collected with scraper and protein lysates were prepared. Protein concentrations were measured using BCA Protein Assay Reagent (Pierce, Rockford, IL). 60µg of each protein sample was loaded to the polyacrylamide gel. Gel was transferred to polyvinylidenedifluoride (PVDF) membrane (TransBlot Turbo, Bio-Rad) with semi-dry transfer according to the manufacturer’s protocol (Bio-Rad, Hercules, CA). Monoclonal anti-ACE2 antibody (ProSci, Poway, CA,1:750diluted in 5% BSA blocking buffer) and goat anti-rabbit IgG-horseradish peroxidase-conjugated antibody (Santa Cruz Biotechnology, Dallas, TX, 1:2500 diluted in 5% BSA blocking buffer) were used as primary and secondary antibodies respectively for ACE2 expression. Monoclonal anti-β-actin antibody (SantaCruz,1:1500diluted in 5%BSA blocking buffer) and goat anti-mouse HRP conjugated (Pierce, Rockford, IL, antibody diluted in 5%BSA blocking buffer) were used as primary and secondary antibodies respectively for β-actin. Membrane images were taken with ChemiDoc MP Imaging System (Bio-Rad, Hercules, CA).

### TMPRSS2 Proteolytic activity assay

The role of Ribavirin in the proteolytic activity of TMPRSS2 was measured both in Vero E6 and Caco-2 cells. To analyze the proteolytic activity of TMPRSS2, 20.000 cells/well were seeded into 96-well plate in 100 µl of DMEM-HG-10%FBS-1%P/S. After 24 hours incubation, the cells were treated with Ribavirin concentrations (5 µM, 10 µM and 25 µM) for 48 hours. Then, the cells were washed with 1xPBS and incubated with fluorogenic synthetic peptide Boc-Gln-Ala-Arg-AMC (Enzo, BML-P237-0005) at a final concentration of 200 uM at 37°Cfor 30minutes. The supernatants were collected separately from each well. They were centrifugated at low-speed (3000 rpm, 10 min at 4°C) to remove the debris. The cleared supernantants were replaced to a new 96 well plate. Fluorescence intensity was measured using fluorescence spectrometer at 380 nm (excitation) and 460 (emission).

### Statistically Analysis

Statistically analysis was performed to evaluate significancy for each data. MTT, Flow cytometry, western blot and proteolytic activity assays were performed in three biological replicates, and qRT-PCR assays were performed in six biological replicates. The median values of the control groups and drug-treated groups were compared with paired two-tailed T test. Analysis of variance was conducted on the replicate values of experiment groups. P values <0.05 was accepted as statistically significant. * indicates that p<0.05. The data was analyzed using Graph Pad Prism 8.01.

## Results

### Ribavirin shows antiviral activity in Vero E6 cells following SARS-CoV-2 infection

Firstly, we performed MTT assay to examine viability in Vero E6 and Caco2 cells after the concentration gradient of the Ribavirin treatment (10, 25, 200, and 750 µM). We observed a significant decrease in the numbers of both cell lines accompanied with increasing concentration of Ribavirin. These data demonstrated that the higher doses (200 and 750 µM) of Ribavirin significantly inhibited the proliferation of these cells in 24 h and 48h post treatment in both cell lines (p < 0.05) (Figure 1A). According to these results, the concentration of 25 µM was assigned to use in the further experiments including antiviral tests since it is the highest concentration while the cell viability is similar to the control group.

**Figure 1.**
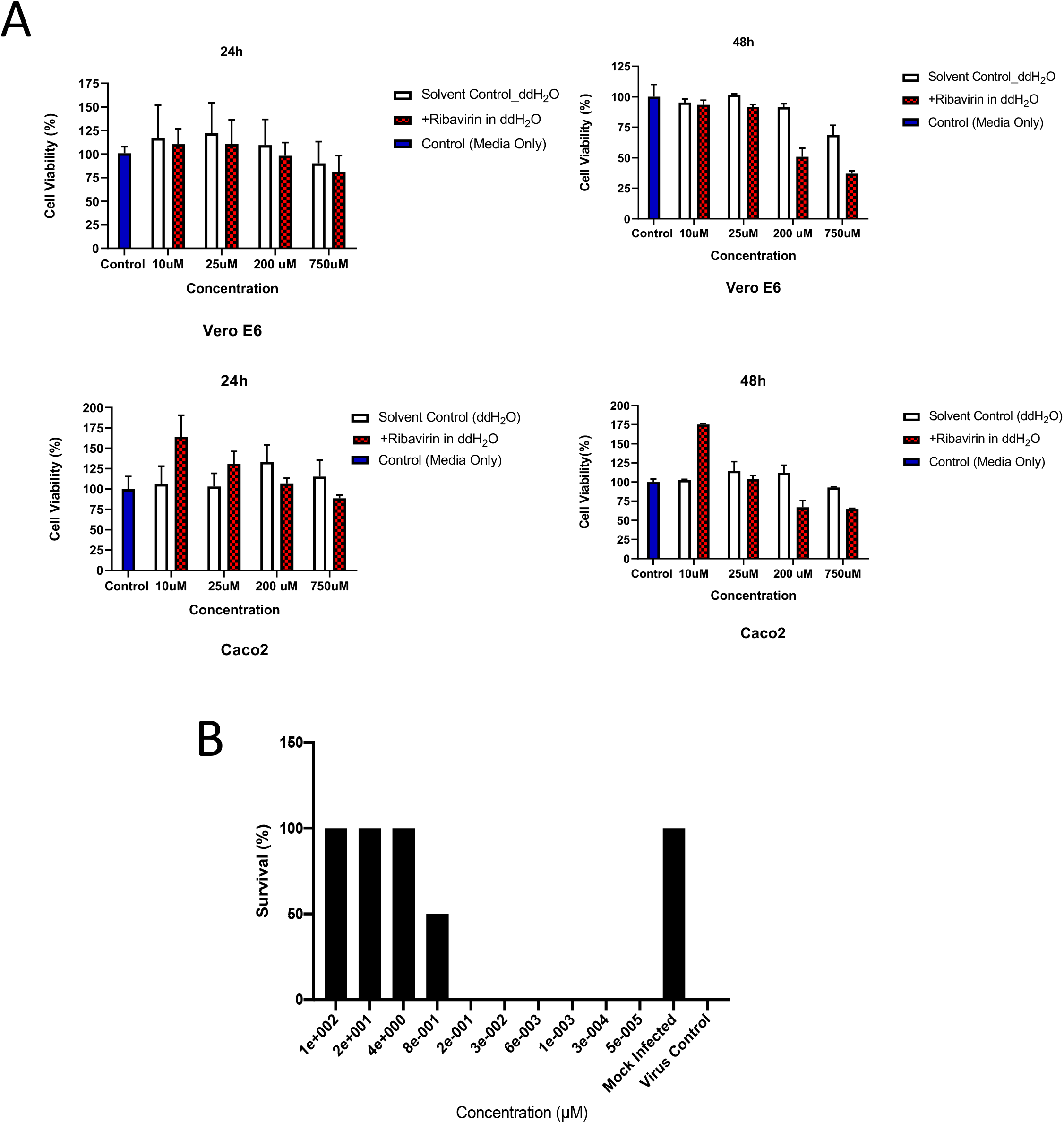
**A)** The effect of Ribavirin at the concentrations of 10, 25, 200, and 750 µM on Vero E6 and Caco2 in 24 and 48 hrs. Data represent the mean and the SD of triplicate samples. Blue bars represent the cells treated with the only cell culture media (Control). White bars represent the cells treated with the cell culture media including DMSO as drug’s solvent (Solvent Control). Red bars represents the cells treated with Ribavirin.. * indicates p<0.05. All data are represented as the mean ± SD (n= 3) **B)** Antiviral activity of ribavirin in vitro. SARS-CoV-2-Ank1 infected Vero E6 cells were treated with Ribavirin drug molecules by serial dilutions. Cells (at 4 different wells) were monitored for CPE using an inverted light microscope. Percentage of cell survival was plotted.

In order to determine the antiviral activity of ribavirin, SARS-CoV-2-Ank1 infected Vero E6 cells were treated with drug molecules across a concentration range (100 µM – 5 pM). Cells at 4 different wells were monitored for CPE using an inverted light microscope. Control cells which were only infected with SARS-CoV-2-Ank1 showed full CPE at day 4. Therefore, the rest of the samples were analyzed at this time point. According to Figure 1B, it was determined that the lowest 4 µM Ribavirin concentration stopped the infectivity of SARS-CoV-2-Ank1, at the level of 100% under *in vitro* conditions. It was also observed that the viral infectivity was stopped at a level of 50% when drug was used at a concentration of 800 nM.

### Ribarivin has a broad-spectrum impact on Vero E6 cells as evident by *in silico* analysis

According to in silico analysis, the affinity energy of Ribavirin toward SARS-CoV-2 proteins and ACE2 is at acceptable values. Also, the affinity of Ribavirin toward different regions of proteins can be accepted as a broad-spectrum impact indicator. Ribavirin interacts with 6M0J (SARS-CoV-2 Spike-ACE2), 6VSB (SARS-CoV-2 Spike Glycoproteins), 6LU7 (COVID19 M-protease), 6M03 (COVID19 M-protease), 6Y84 M^pro^, 6LXT (Fusion Protein), 5×29 (envelope (E) protein), 6M71 (RNA polymerase), 6VWW (Nsp15 Endoribonuclease), 6VYO (RNA binding region of Nucleocapsid Protein), and 1R42 (ACE2) proteins, and affinity energies of Ribavirin change in the range between -7.5 and -3.3 kcal/mol (Figure 2). Abundance of H-bonds and Ribavirin acting as both hydrogen donor and acceptor increase the binding strength of Ribavirin to the protein surface. On the other hand, SAS calculations of SARS-CoV-2 proteins showed high binding (green field) and high SAS value (blue field) for Ribavirin affected regions (Figure S1, S2 and S3). These indicate constructed H-bonds by Ribavirin are exposed to the strong impact of H_*2*_O molecules. Therefore, the surface attachment time of Ribavirin is affected.

**Figure 2:**
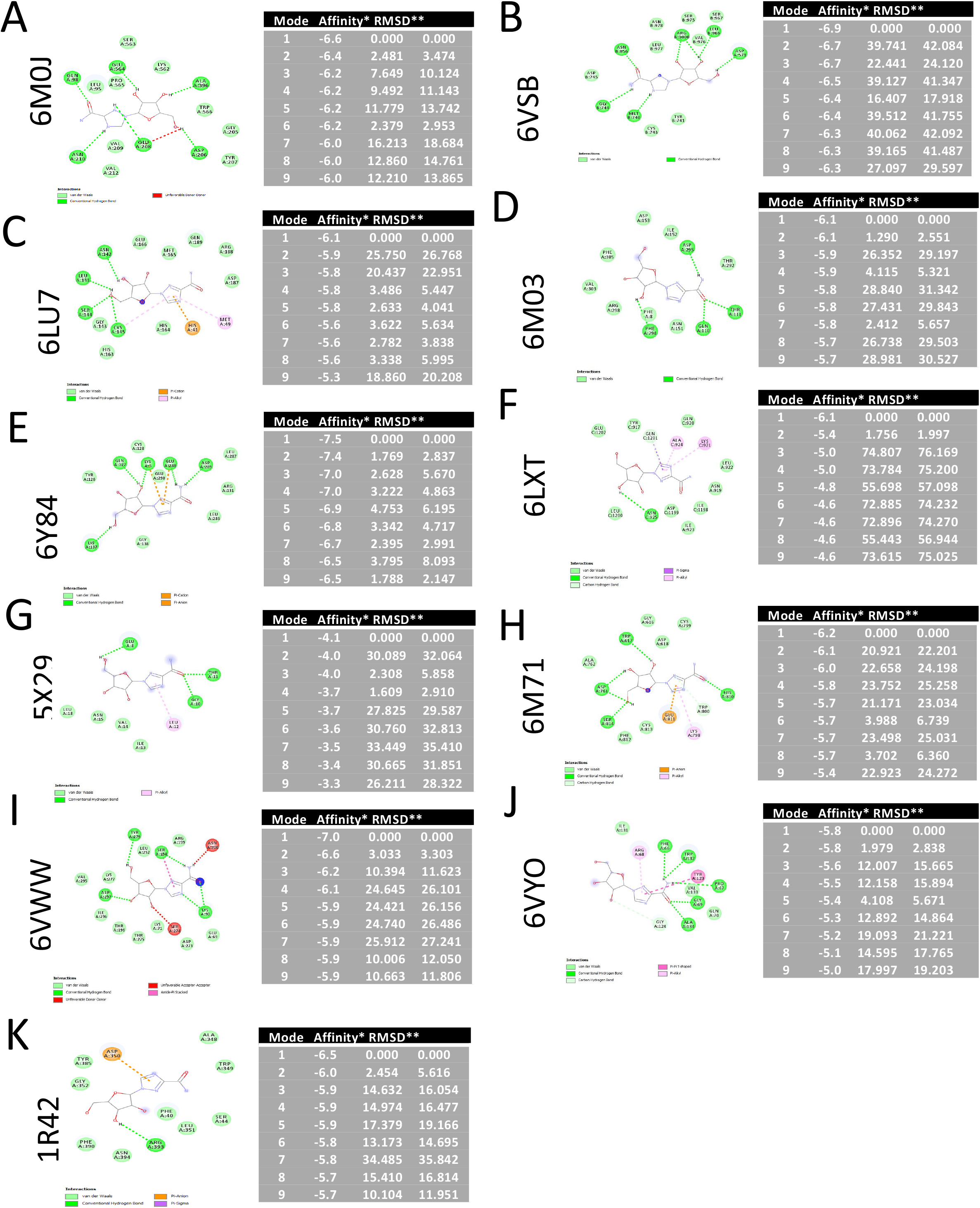
2D representation and docking scores of ribavirin and protein interaction. 6M0J: SARS-CoV-2 Spike ACE2, 6VSB: SARS-CoV-2 Spike Glycoproteins, 6LU7: COVID19 M-protease, 6M03: COVID19 M-protease, 6Y84: M^pro^, 6LXT: Fusion Protein, 5×29: envelope (E) protein, 6M71: RNA polymerase, 6VWW: Nsp15 Endoribonuclease, 6VYO: RNA binding region of Nucleocapsid Protein, 1R42: ACE2 proteins

### Ribavirin decreases TMPRSS2 expression in Vero E6 cells

Since in silico docking analysis showed that Ribavirin has a broad-spectrum impact on Vero E6, we performed further molecular analysis to delineate the effect of the drug on viral lifecycle. TMPRSS2 enables the virus entry [13], therefore we aimed to test whether Ribavirin has an inhibitory effect on TMPRSS2 and ACE2. Firstly, we tested mRNA expression levels of TMPRSS2 in the25 µM Ribavirin-treated Vero E6 and Caco 2 cells by qRT-PCR. Caco 2 cells were included in this study to show the effectivity in human cell line other than Vero E6. We demonstrated that the Ribavirin treatment significantly leads to a two fold decrease comparing to solvent control in the TMPRSS2 expression in 48 hours in the Vero E6 cells (Figure 3A). On the contrary, any significant change was not observed at 48th hours in Caco 2 cells (Figure 3A). To address the question whether the TMPRSS2 protein expression level is altered with the Ribavirin treatment, we performed flow cytometry assay in the Vero E6 and Caco-2 cells following 25 µM Ribavirin treatment for 48 hours. TMPRSS2 level was significantly decreased in Ribavirin treated group comparing to the control. (For Vero-E6; Control cells 77.6 ± 14.55 %, Solvent treated cells 70.65 ± 6.5%, Ribavirin treated cells 40.89 ± 21.9%; for Caco2; Control cells 52.15 ± 3.18%, Solvent treated cells 56.43 ± 0.77%, ribavirin treated cells 34.2 ± 3.05%) (Figure 3B). These data suggested that Ribavirin treatment significantly reduces TMPRSS2 expression both at protein and gene level at the end of the 48 hours-treatment.

**Figure 3:**
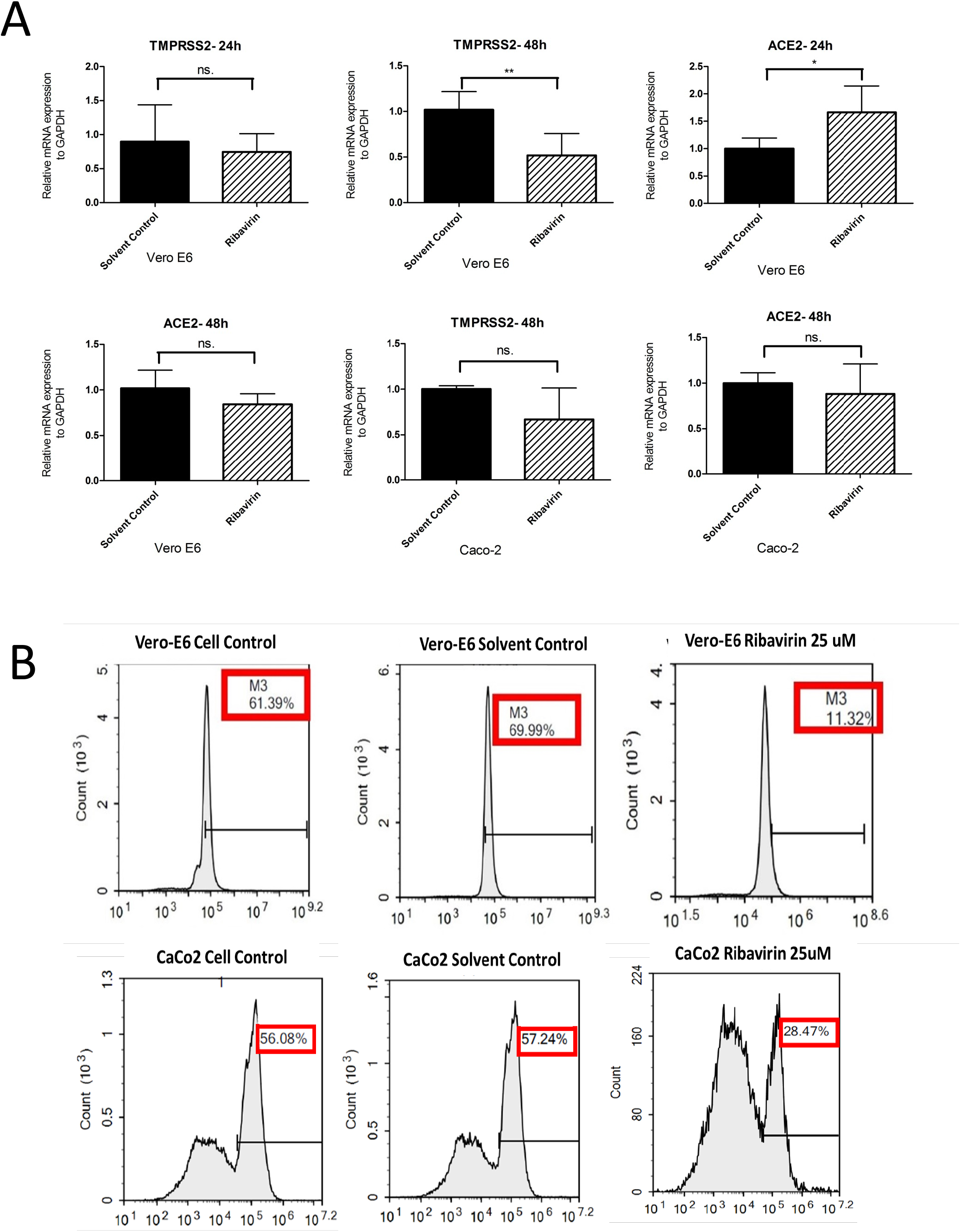
Ribavirin decreases TMPRSS2 expression in Vero E6 cells. qRT-PCR demonstrates relative fold differences of TMPRSS2 mRNA expression after 24 h and 48 h Ribavirin treatment in **A)** Vero-E6 cells and **B)** Caco 2 cells. (P<0.05). * indicates p<0.05. All data are represented as the mean ± SD (n= 6). **C)** Representative figure shows flow cytometry analysis of the Ribavirin (25 µM) treated Vero-E6 and Caco2 cells. The Ribavirin treated cells indicates a significant decrease in the TMPRSS2 expression both in Vero-E6 and Caco2 cells (Paired Two-tailed T test, P<0.05). * indicates p<0.05. All data are represented as the mean ± SD (n= 3).

### The suppressive effect of Ribavirin in ACE2 protein expression changes depending on the cell type

In addition to evaluation of protein level of TMPRSS2, to elucidate the mechanism of Ribavirin in the regulation of ACE2 expression at protein level we performed western blotting assay. Our qRT-PCR studies have demonstrated that Ribavirin treatment does not any effect on mRNA level of ACE2 at 48th hours both in Vero-E6 and Caco2 cells. Consistently with this result, we observed the there is no significant effect of Ribavirin in the regulation of protein level of ACE2 in in Vero-E6. On the other hand, in contrast with mRNA expression results of Caco2 cells, a significant reduction in ACE2 protein level was detected at 48th hours-treatment (Figure 4).

**Figure 4:**
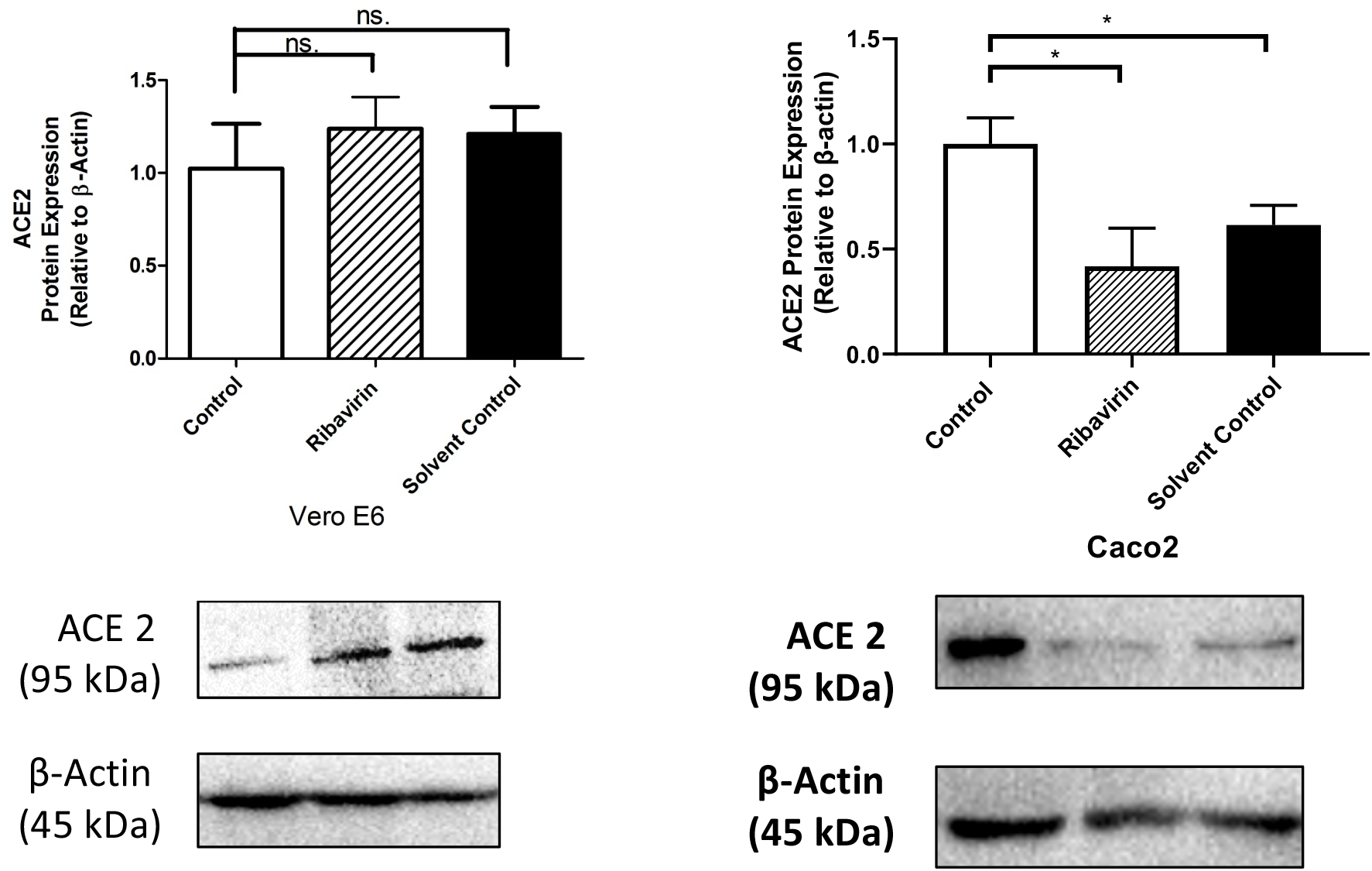
The suppressive effect of Ribavirin in ACE2 protein expression. Representative bands and analysis of band intensities show protein changes of ACE2 in Ribavirin (25 µM) treated Vero-E6 and Caco2 cells. The Ribavirin treated Caco-2 cells indicates a significant decrease in the ACE2 expression. But ribavirin treatment does not cause any significant change in Vero E6 cells. Solvent treatment was used as a negative control. * indicates p<0.05. All data are represented as the mean ± SD (n= 3).

### Ribavirin has an inhibitory effect in proteolytic activity of TMPRSS2 enzyme

A well-known mechanism is that SARS-CoV-2 entries a host cell using its spike protein appearing on the surface of the virus binds ACE2 receptor on the host cell surface. After this binding, the virus primes two cut sites and causes a conformational change which allows the fusion of viral and host membranes. Besides ACE2, TMPRSS2 has an extracellular protease domain that cleaves the spike protein to begin membrane fusion. Due to this reason the inhibition of TMPRSS2 protease activity might be a valuable and important sign to analyze the anti-viral effect of drug at molecular level [14]. *In vitro* measurement of a fluorescence resonance energy transfer of a synthetic protease substrate (Boc-Gln-Ala-Arg-AMC, ENZO Life Sciences) is a very useful tool to examine the TMPRSS2-activated matriptase proteolytic activities [15]. The substrate protein was added into Vero-E6 and Caco-2 cell cultures which were treated with 5, 10, and 25 µM Ribavirin for 48 hours. In accordance with the measurement of soluble forms of proteins released from cells, we detected proteolytic activity of TMPRSS2 in the supernatant of cell culture. It was observed that Ribavirin treatment significantly reduces TMRPRSS2 activity in a dose dependent manner both in Vero-E6 and Caco-2 (Figure 5) cells comparing to the control. This data indicates that Ribavirin has an inhibitory role in the proteolytic activity of TMPRSS2 enzyme.

**Figure 5:**
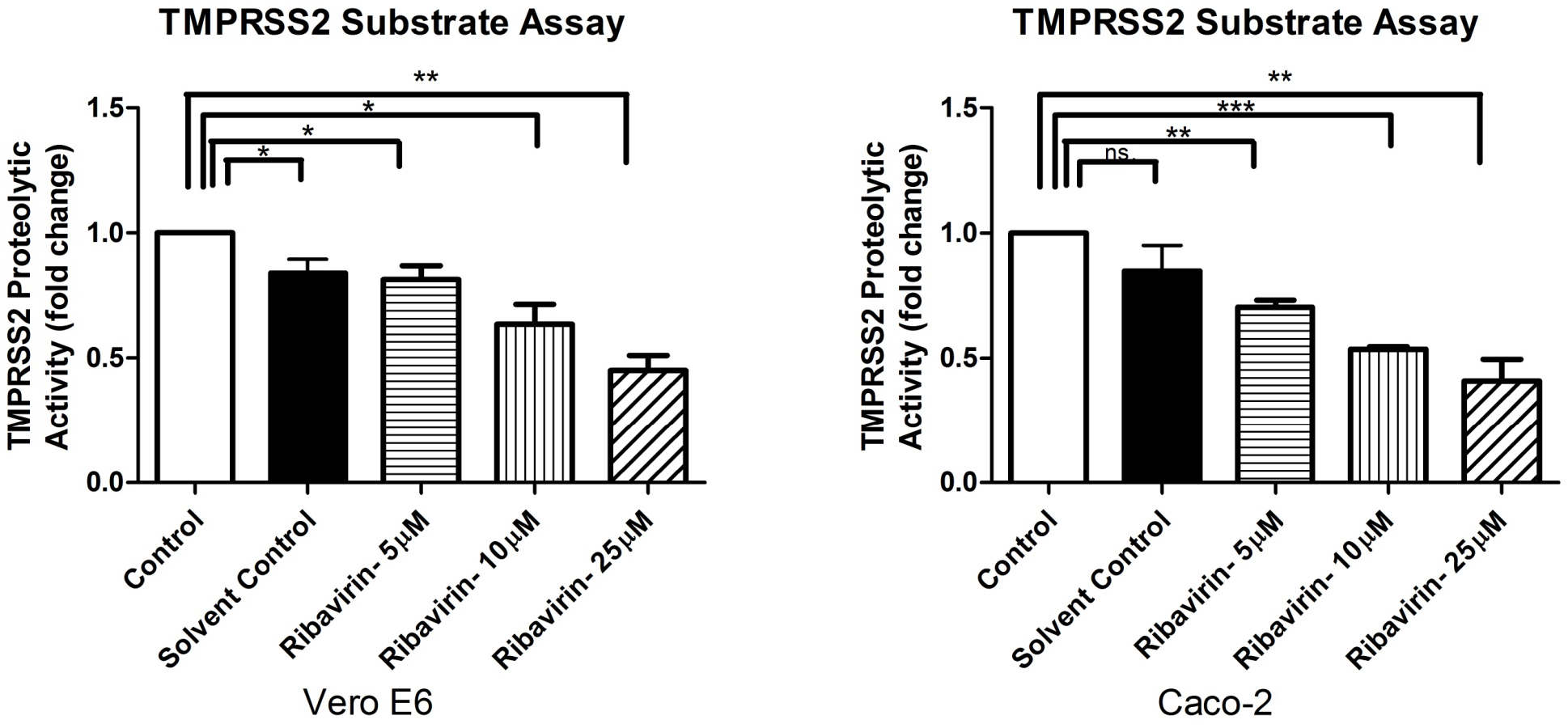
Evaluation of proteolytic activity of TMPRSS2 enzyme was performed via substrate assay analysis. In accordance with the measurement of soluble forms of proteins released from cells, we detected proteolytic activity of TMPRSS2. Ribavirin treatment causes significantly decrease of primed substrate by TMPRSS2. This data indicates that Ribavirin suppresses proteolytic activity of TMPRSS2 enzyme. Solvent treatment was used as a negative control. * indicates p<0.05. All data are represented as the mean ± SD (n= 3)

**Figure 6:**
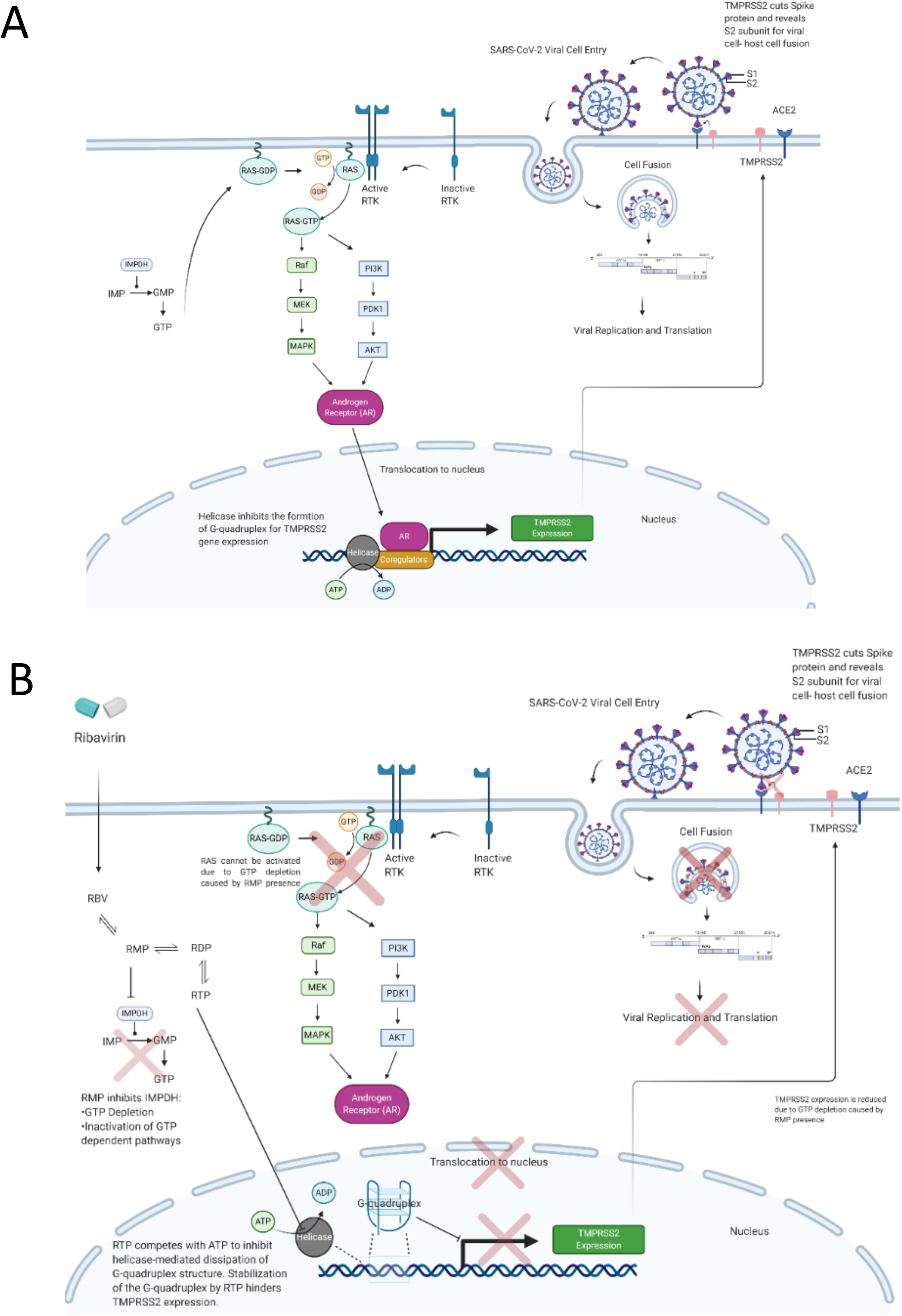
SAR-CoV-2 viral entry and TMPRSS2 expression influenced by Ribavirin treatment. **A)** Spike protein on SARS-CoV-2 virus interacts with angiotensin converting enzyme 2 (ACE2) on host membrane. Transmembrane serine protease 2 (TMPRSS2) located on host membrane cleaves the SARS-CoV-2 Spike protein and releases S2 subunit. S2 subunit induces cell fusion between host cell and SARS-CoV-2. After cell fusion positive single strand viral RNA (+ssRNA) exposed in the host cytosol and uncoating of +ssRNA induces viral replication and translation. TMPRSS2 expression is driven by RAS-GTP dependent androgen receptor (AR) signaling. GTP is produced through inosine monophosphate (IMP) catalyed with ionisine monophosphate dehydrogenase (IMPDH) enzyme. Also, the upstream of TMPRSS2 gene promoter contains G-rich sequence which creates tendency to create G-quadruplex formation that inhibits gene expression. Helicase protein inhibits the G-quaruplex formation. **B)** Ribavirin enters cells via cell membrane and turns into ribavirin monophosphate (RMP), ribavirin diphosphate (RDP) and ribavirin triphosphate (RTP) by phosphorylation. RMP inhibits IMPDH enzyme activity and causes GTP depletion in the cell. GTP depletion leads to inhibition of activation of RAS-GDP complex. If the RAS-GDP complex cannot be activated with GTP downsignalling AR pathway cannot be activated and cannot induce TMPRSS2 expression which is leading to decrease in TMPRSS2 expression. RTP competes with ATP thereby, inhibits the helicase activity in the nucleus leading stabilization of G-quadruplex complex on upstream of TMPRSS2 gene promoter to inhibit TMPRSS2 gene expression. Decreased expression of TMPRSS2 protein induces inhibition of viral cell entry.

## Discussion

Ribavirin is a guanosine analog and has a broad-spectrum antiviral activity against RNA viruses. Based on this, we aimed to show the anti-SARS-CoV-2 activity of this drug molecule via in vitro, in silico and molecular techniques. In vitro inhibition of viral infection suggested that Ribavirin has antiviral activity when used at a concentration above 800nm. Later, in silico analysis of the interaction between Ribavirin and 9 possible domains involved in SARS-CoV-2 infection was performed. Results suggested that although favorable binding affinities were observed between drug molecules and protein domains, the mechanism of action could not be explained via a single domain interaction.

SARS-CoV-2 virus is composed of a positive sense single stranded RNA (+ssRNA) genome, spike (S), envelope (E), membrane (M), and nucleocapsid (N) proteins. S protein composed of S1 and S2 domains. S1 domain of S protein contains receptor-binding domain (RBD) and interacts with angiotensin converting enzyme 2 (ACE2) receptor located on host organism membrane for viral cell entry [16, 17]. Transmembrane protease serine 2 (TMPRSS2) enzyme located on host cell membrane cleaves S protein and reveals S2 domain of S protein which leading to fusion of the viral membrane with the host cell membrane to release viral +ssRNA genome into a host cell cytoplasm [18, 19]. Chloroquine phosphate and hydroxychloroquine suggested to inhibit ACE2 [20]. Camostat mesylate and Nafamostat mesylate suggested for the inhibition of TMPRSS2 priming [21]. Therefore, TMPRSS2 is one of the potential targets for anti-SARS-CoV-2 drug development. In this study, we have shown that Ribavirin treatment of Vero E6 cells decreased the expression of TMPRSS2 which could explain the mechanism of action against SARS-CoV-2. The mechanism linking ribavirin to TMPRSS2 expression is beyond the scope of the present study, however we have identified a few possible mechanisms. A possible explanation for the observed decrease in TMPRSS2 expression is that ribavirin-mediated GTP depletion hinders the signaling cascades that activate the androgen receptor (AR), a potent activator of TMPRSS2 expression [22, 23]. Ribavirin monophosphate (RMP)-mediated inhibition of inosine monophosphate dehydrogenase (IMPDH) is a prominent antiviral mechanism occurring even at relatively low concentrations of ribavirin [22, 23]. IMPDH is responsible for the production of GMP (a GTP precursor) from inosine monophosphate (IMP). RMP competes with IMP for binding to IMPDH, thereby reducing GMP and GTP synthesis [24]. Competitive inhibition of IMPDH by RMP depletes local GTP, causing downstream imbalance of nucleotides that interferes with mRNA capping, ribosomal function, and G-protein signaling [22, 24]. Because AR is activated by the MAPK pathway involving the G-protein Ras [25], one could conjecture that depletion of GTP could indirectly dampen AR activity and by extension TMPRSS2 expression. Ribavirin-mediated depression of Ras-signaling has not been demonstrated experimentally, but the theoretical connection is worth future exploration.

Ribavirin may also inhibit the expression of TMPRSS2 by modulating the formation of inhibitory G-quadruplex structures at the TMPRSS2 promoter. In a G-quadruplex, G-rich sequences form tetrads around K+ ions and stack to form the G-quadruplex structure [26, 27]. Formation of G-quadruplex structures in the promoter region can influence gene expression, and some cancer therapeutics have been explored for stabilizing these structures to silence proto-oncogenes [28]. A recent study identified a guanosine rich sequence upstream of the TMPRSS2 promoter, capable of forming a G-quadruplex structure [29]. Stabilization of G-quadruplex in the TMPRSS2 promoter inhibited TMPRSS2 expression and IAV replication [29]. G-quadruplexes can be unwound by helicase enzymes to relieve the suppression of downstream genes. Ribavirin triphosphate inhibits the activity of helicases by competitive inhibition of ATP-hydrolysis [30]. We propose that ribavirin-mediated G-quadruplex stabilization could potentially explain the observed decrease in TMPRSS2 expression.

## Acknowledgement

Authors would like to acknowledge the funding from the Scientific and Technological Research Council of Turkey (TUBITAK) under the grant number 18AG020.

