## supplemental files for "Ribavirin shows antiviral activity against SARS-CoV-2 and downregulates the activity of TMPRSS2 and the expression of ACE2 In Vitro"

### Supplementary Figures

6M0J SARS-CoV-2 Spike ACE2

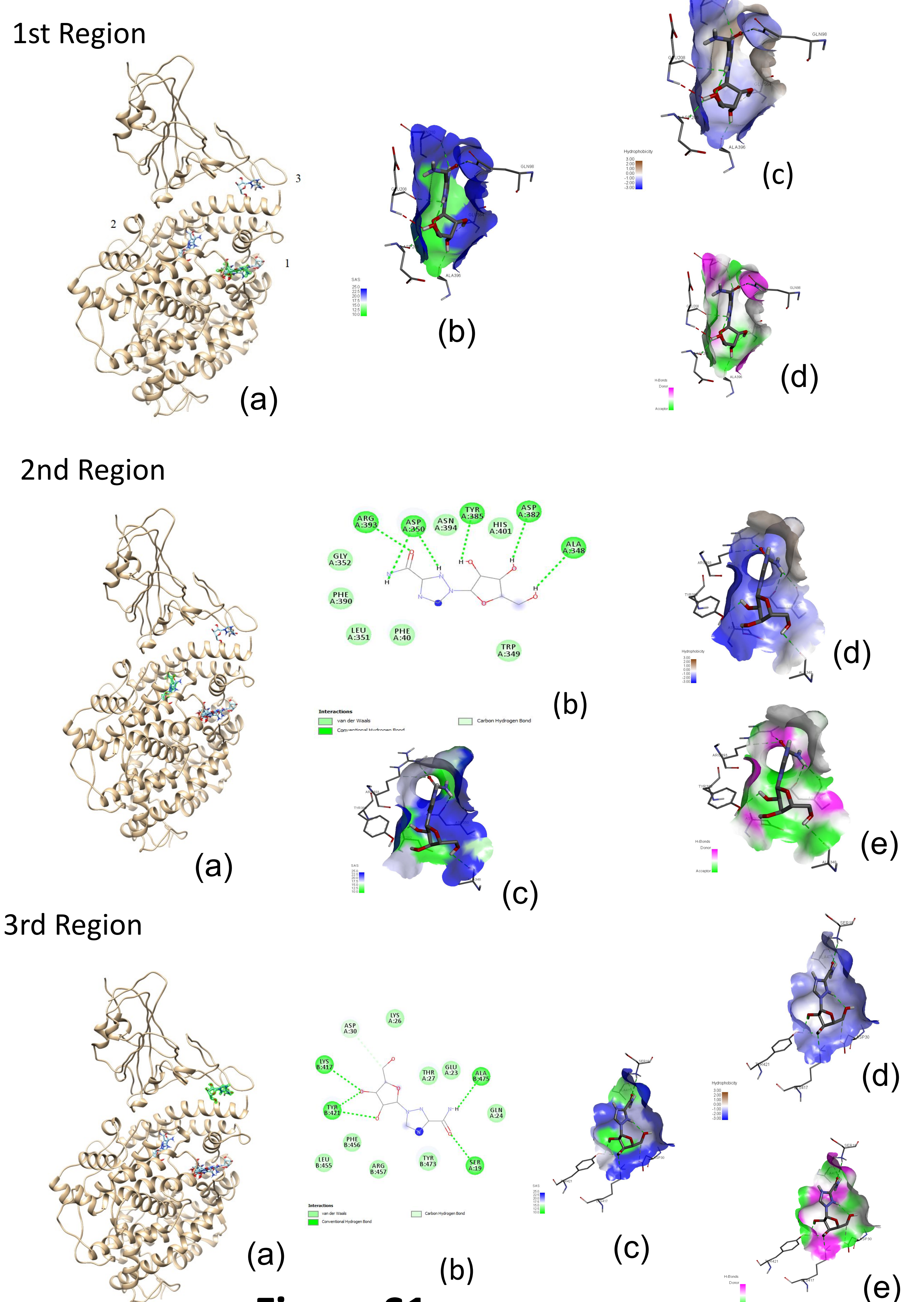


**Figure S1**

SARS CoV-2 Spike glycoproteins: 6VSB (open form) and 6VXX (close form)

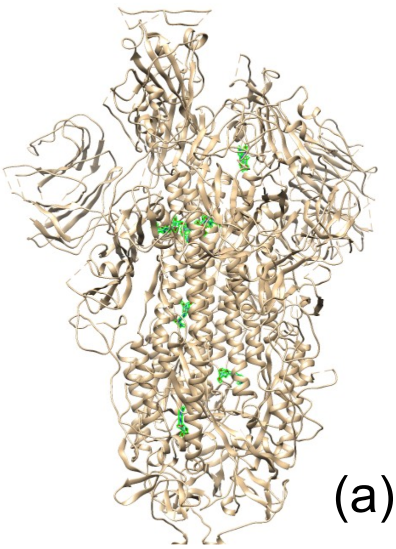

(a)

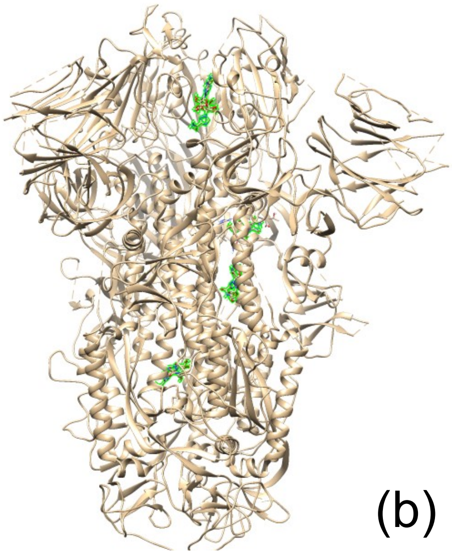

(b)

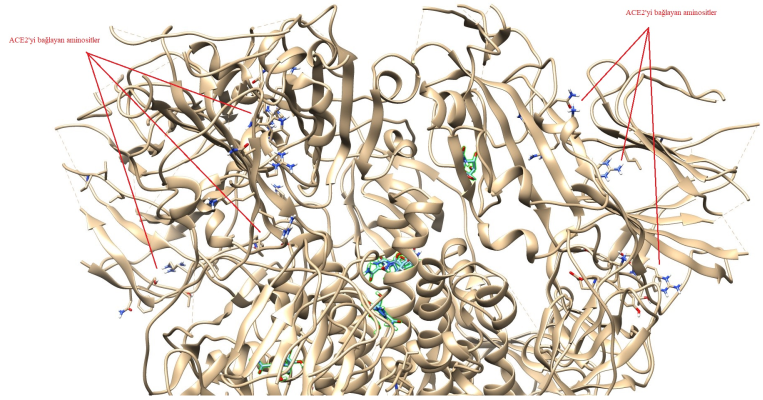

c)

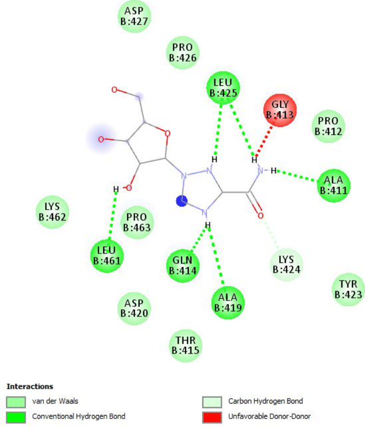


5th conformation

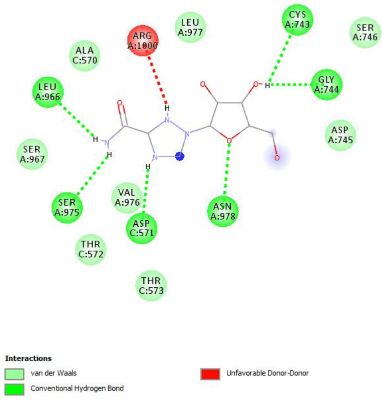


6th conformation

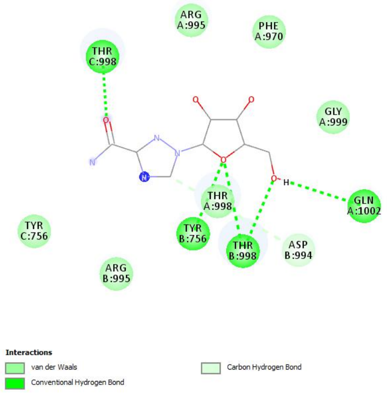


7th conformation

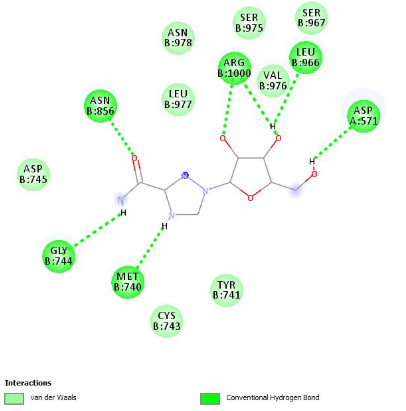


8th conformation

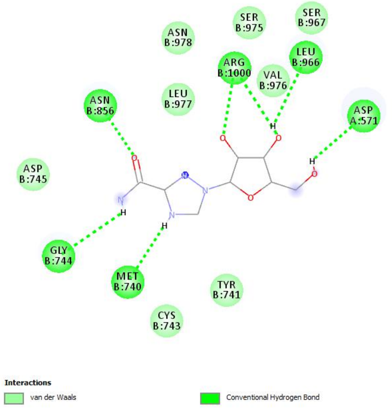


9th conformation

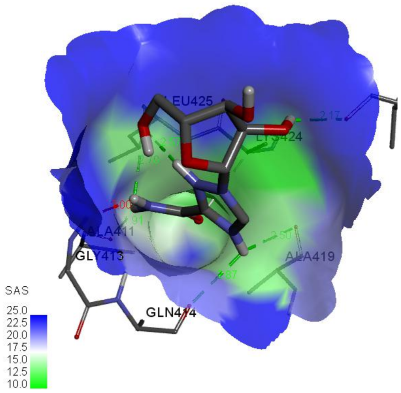


Eligibility of solvent in a region where 5th conformation located (SAS)

### Figure S2

#### 6LU7 COVID-19 M-protease

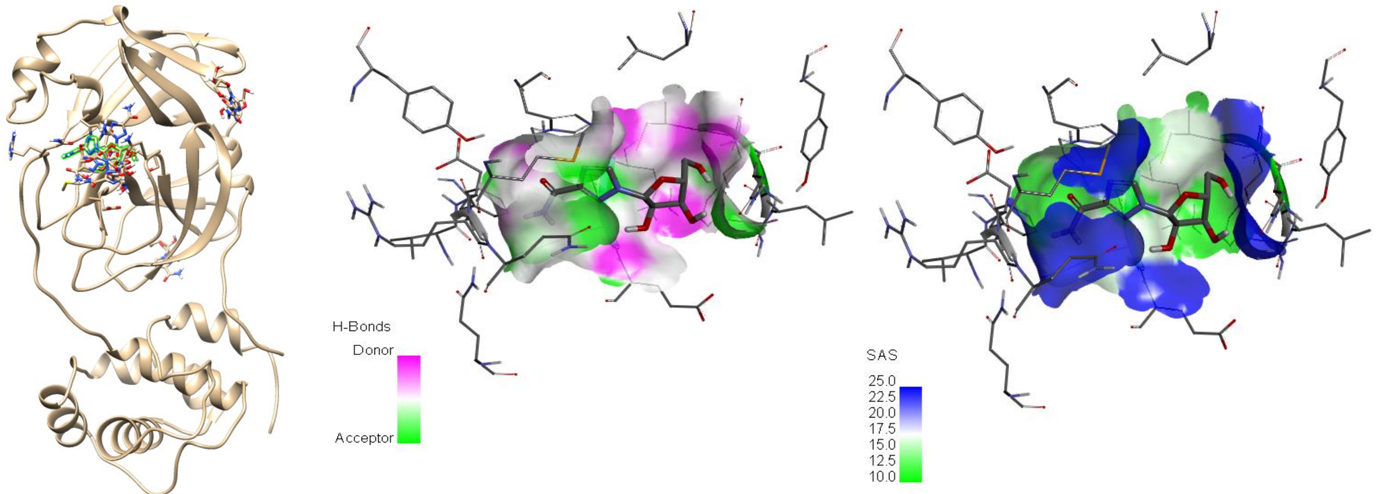


(a) (b) (c)

#### 6M03 COVID-19 M-protease

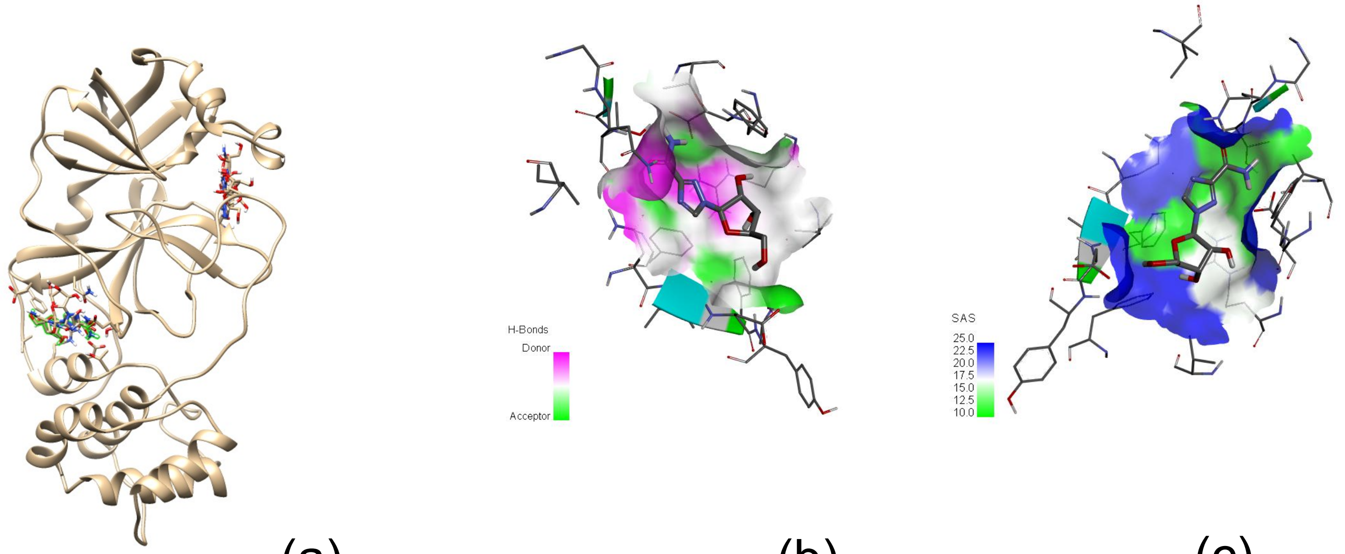


(a) (b) (c)

#### 6Y84 COVID-19 M-protease

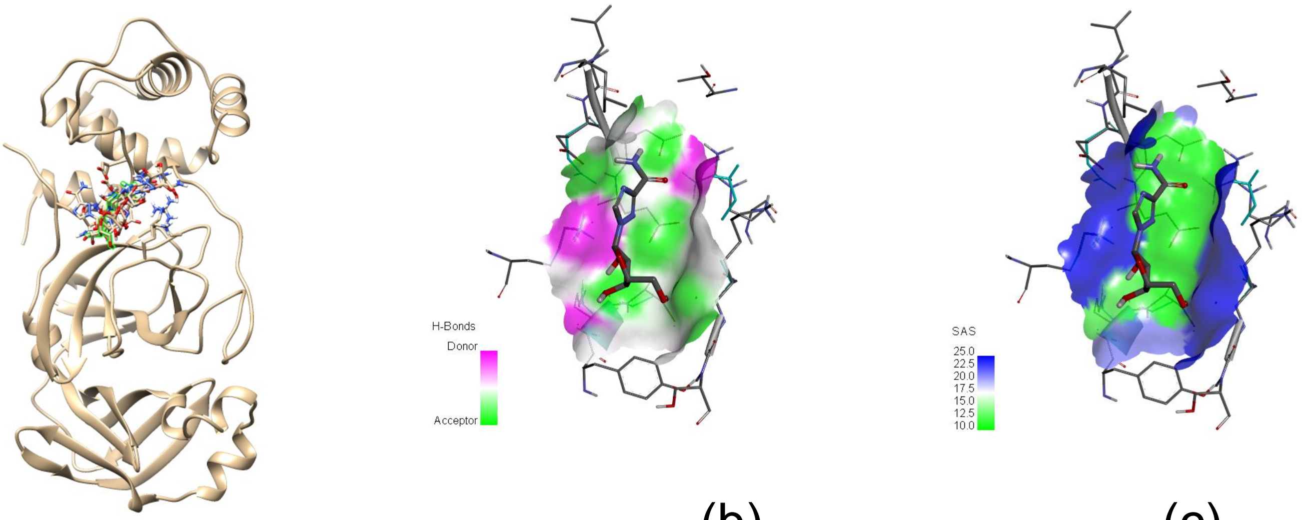


(a) (b) (c)

### Figure S3
